# Drosophila immune priming to Enterococcus faecalis relies on immune tolerance rather than resistance

**DOI:** 10.1101/2022.07.20.500468

**Authors:** Kevin Cabrera, Duncan S. Hoard, Daniel I. Martinez, Zeba Wunderlich

## Abstract

Most multicellular organisms, including fruit flies, possess an innate immune response, but lack an adaptive immune response. Even without adaptive immunity, “immune priming” allows organisms to survive a second infection more effectively after an initial, non-lethal infection. We used *Drosophila melanogaster* to study the transcriptional program that underlies priming. Using an insect-derived strain of Gram-positive *Enterococcus faecalis*, we found a low dose infection enhances survival of a subsequent high dose infection. The enhanced survival in primed animals does not correlate with a decreased bacterial load, implying that the organisms tolerate, rather than resist the infection. We measured the transcriptome associated with immune priming in the fly immune organs: the fat body and hemocytes. We found many genes that were only upregulated in re-infected flies. In contrast, there are very few genes that either remained transcriptionally active throughout the experiment or more efficiently re-activated upon reinfection. Measurements of priming in immune deficient mutants revealed IMD signaling is largely dispensable for responding to a single infection, but needed to fully prime; while Toll signaling is required to respond to a single infection, but dispensable for priming. Overall, we found a primed immune response to *E. faecalis* relies on immune tolerance rather than bacterial resistance and drives a unique transcriptional response.

## Introduction

The fruit fly *Drosophila melanogaster* inhabits environments rich in bacteria, fungi, and viruses. The fly has to mitigate these pathogens to survive. To this end, it has evolved a tightly controlled innate immune response. It has long been appreciated that the fly immune pathways can distinguish between Gram-positive bacteria and fungi versus Gram-negative bacteria (Buchon, et al. 2014). Recent findings have elaborated on these models by showing specificity within Gram-classifications, cross-talk between the two individual pathways, and a remarkable level of additional molecular coordination (Kleino, et al. 2014; Lin, et al. 2020; Hanson, et al. 2019).

Among these refined characteristics is the potential for immune memory in the innate immune system. While flies lack the canonical antibody-mediated immune memory of the adaptive immune response, an initial non-lethal infection can sometimes promote survival of a subsequent infection. This phenomenon, termed immune priming, has been observed in evolutionarily distant organisms such as plants (Cooper & Ton 2022), multiple arthropod species (Milutinović, et al. 2016), and mammals (Netea, et al. 2016; Divangahi, et al. 2020). The fact that this mechanism is present in animals that have an adaptive response hints at its importance in organismal fitness.

Despite immune priming’s effect on survival, the underlying mechanism controlling it in flies is not completely understood. Three mechanistic hypotheses have been proposed to explain the physiological effects of priming (Cooper & Eleftherianos 2017; Coutasu, Kurtz, Moret 2016). The first is that there is a qualitatively different response in how primed insects react to an infection versus non-primed insects, leading to a more effective response. A second hypothesis is that insects will initiate an immune response during priming, but will re-initiate the same immune function in a potentiated manner upon reinfection. This is most similar to the phenomenon of what has been observed in mammalian trained immunity (Divangahi, et al 2020). Lastly, immune effectors created during the initial immune response may loiter in the body, eliminating the lag time in initiating effector production. Since flies often harbor low-level chronic infections instead of completely clearing them (Duneau, et al. 2017; Chambers, et al. 2019), these chronic infections may contribute to immune priming by providing a consistent mild stimulus. However, it could be that priming is driven by a combination of these three mechanisms. Delineating the relative contributions of each of these mechanisms may not only reveal the drivers of infection survival, but may also suggest epigenetic mechanisms of gene regulation and tradeoffs between the immune response and other biological processes.

*Drosophila* is a good model for dissecting the mechanisms driving immune priming due to its genetic tractability, extensively characterized innate immune pathways, and its homology to mammalian innate immune pathways. There has been extensive characterization of the fly’s transcriptional response to a variety of bacteria (Troha, et al. 2018; Schlamp, et al. 2021; De Gregorio, et al. 2002) and the progression of bacterial load during infection with different bacteria or in different host genotypes (Duneau, et al. 2017). Studies of priming have revealed the key role of phagocytosis. Blocking phagocytosis in adults decreases priming with the Gram-positive bacterium *Streptococcus pneumoniae* (Pham, et al. 2007). Blocking developmental phagocytosis of apoptotic debris also makes larvae more susceptible to bacterial infection (Weavers, et al. 2016). In addition, the production of reactive oxygen species as a result of wounding contributes to immune priming with the Gram-positive bacterium *Enterococcus faecalis* (Chakrabarti & Visweswariah 2020). These findings lay the foundation for testing the mechanistic hypotheses that underlie immune priming.

In this study, we present a multifaceted approach to understand immune priming in the fly using an *E. faecalis* reinfection model. *E. faecalis*, a Gram-positive, naturally occurring pathogen of the fly, has been previously used to induce an immune response with dose-dependent lethality. We characterize not only the physiological response to priming by way of survival and bacterial load to immune priming, but also the transcriptional response that underlies the physiology. By assaying transcription separately in both the hemocytes and fat body, we explore the organ-specific program that mounts a more effective primed immune response.

## Results

### *E. faecalis* priming increases survival after re-infection

To determine whether we could elicit a priming response in flies, we needed to find appropriate priming and lethal doses. For these experiments, 4-day old male Oregon-R flies were infected with a strain of the Gram-positive bacteria *Enterococcus faecalis* originally isolated from wild-caught *D. melanogaster* (Figure 1A) (Lazarro, et al. 2006). Initial infection with *E. faecalis* showed dose-dependent survival (Figure 1B). Flies infected with a dose of ∼30,000 CFU/fly (*Efae* High Dose) gradually died off, with more than fifty percent of flies dying by day 2, making it a practical choice for representing a lethal dose. Flies injected with a lower dose of ∼3,000 CFU/fly (*Efae* Low Dose) had survival comparable to those injected with PBS, indicating that death was largely due to the injection process itself, rather than from bacterial challenge.

**Figure 1.**
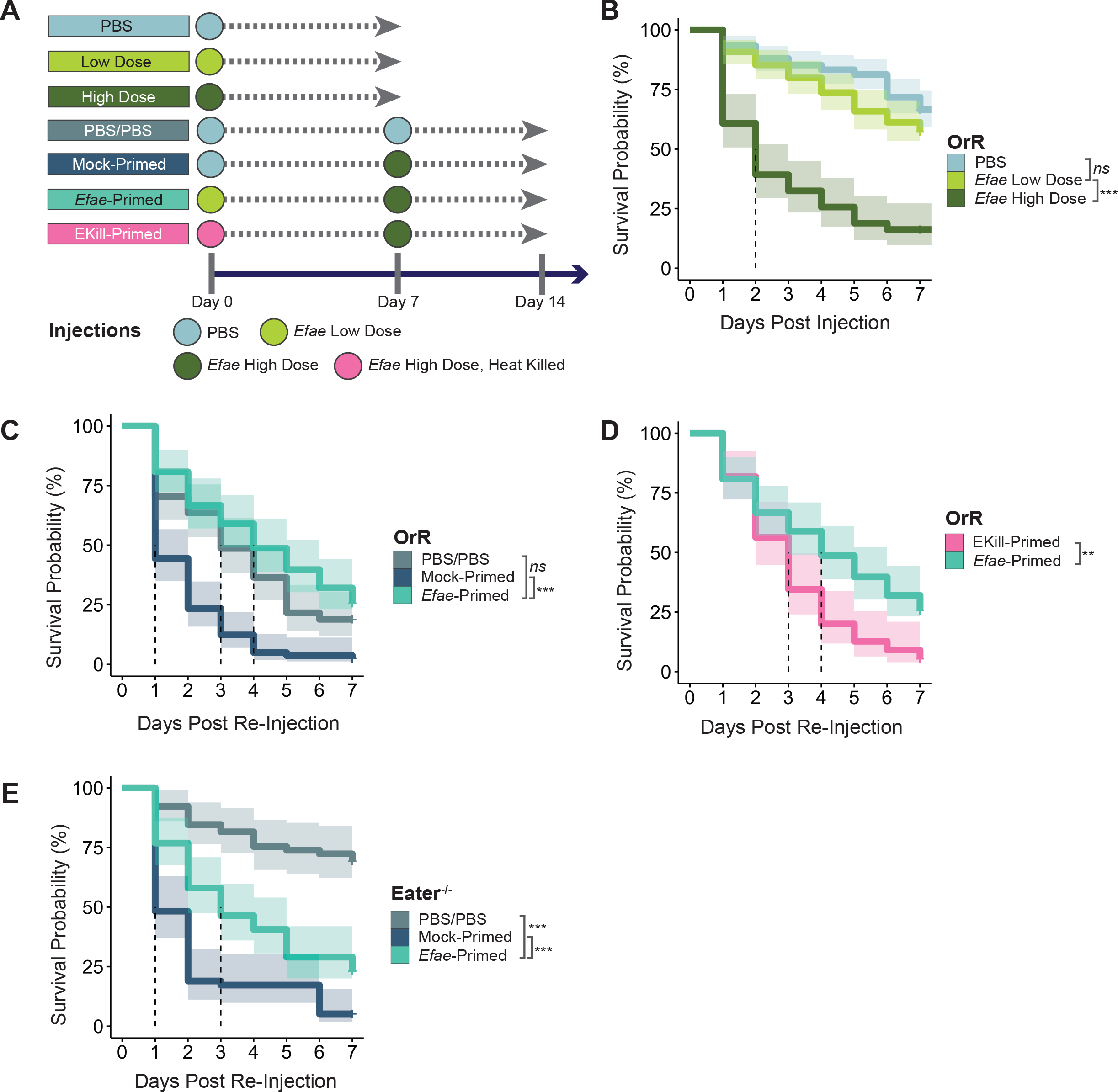
*E. faecalis* can induce immune priming in *D. melanogaster*. **A)**. Schematic of single and double-injection experiments. **B)**. Survival of Oregon-R flies injected with PBS (n = 149), *Efae* Low Dose (∼3,000 CFU/fly, n = 129), and *Efae* High Dose (∼30,000 CFU/fly, n = 74). Dotted line indicates median survival time. Shaded area indicates 95% confidence interval. PBS vs Low Dose: p = 0.081; Low Dose vs. High Dose: p < 0.0001; all survival significance testing is log rank-sum test [* p<0.01, ** p<0.001, *** p<0.0001] **C)**. Survival of primed OrR flies versus double-injected, non-primed controls (PBS/PBS: n = 74, Mock-Primed: n = 81, *Efae*-Primed: n=78). PBS/PBS vs *Efae*-Primed: p = 0.13; Mock-Primed vs. *Efae*-Primed: p < 0.0001 **D)**. Survival of OrR flies primed with heat-killed *E*.*faecalis* (EKill-Primed: n = 55) versus flies primed with live *E*.*faecalis*: p = 0.00068. **E)**. Survival of primed phagocytosis-deficient, *eater*-mutant flies versus double-injected, non-primed controls (PBS/PBS: n = 65, Mock-Primed: n = 58, *Efae*-Primed: n=69). PBS/PBS vs *Efae*-Primed: p < 0.0001; Mock-Primed vs. *Efae*-Primed: p < 0.0001.

To model re-infection, flies were initially injected either with a low bacterial dose (i.e. *Efae*-primed flies) or a negative control of PBS (i.e. Mock-primed flies) (Figure 1A). After resting for seven days, flies were re-injected with a high dose of *E. faecalis* and assayed. Seven days was chosen as the priming interval because we found that flies had gained enhanced re-infection survival from priming (Supplementary Figure 1A), reached a stable chronic bacterial load (Figure 2A), and survived in high enough numbers to practically collect for re-infection. The median survival time after re-injection was significantly increased from Mock-primed flies (1 day) to *Efae*-primed flies (4 days) (Figure 1C). Though there was a decrease in survival from double wounding compared to a single wound (Supplementary Figure 1B), *Efae*-primed flies still had greater survival compared to this double-injected baseline as well as when compared to single, High Dose-infected flies (Supplementary Figure 1C). Priming with heat-killed *E. faecalis*, which retains its signaling-responsive components but lacks any additional virulence factors (Itoh, et al. 2012; Adams, et al. 2010), resulted in a more moderate increase in survival rate compared to live bacteria priming (Figure 1D). This implies some level of priming is conferred simply through bacterial sensing, but that the effect is not as robust as when the fly is exposed to the live microbe.

**Figure 2.**
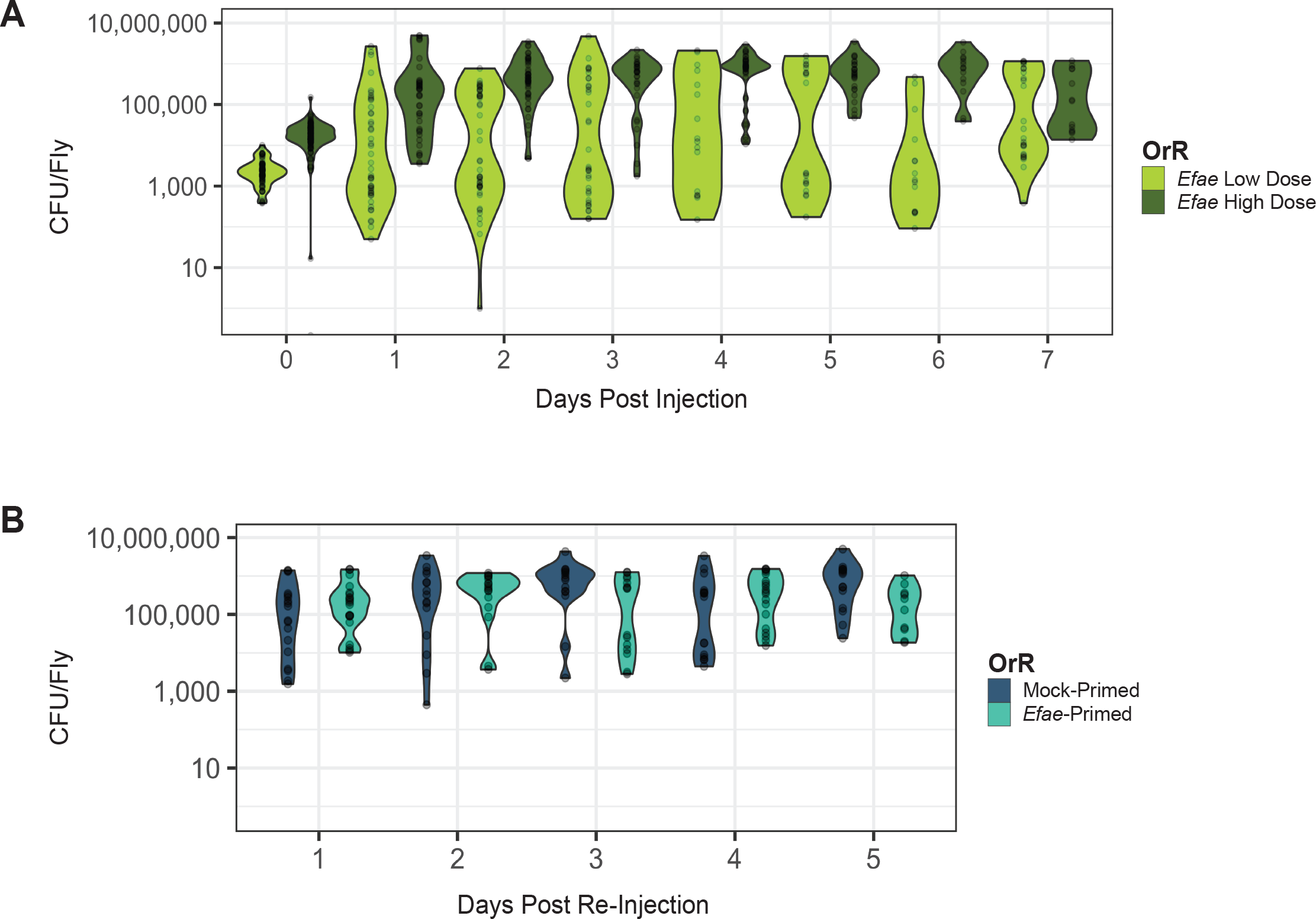
Bacterial clearance is not correlated with primed survival against *E. faecalis* re-infection. **A)**. Bacterial load of single-injected flies. Flies were abdominally injected with either *E*.*faecalis* Low Dose (∼3,000 CFU/fly) or *E. faecalis* High Dose (∼30,000 CFU/fly), and a subset was dilution plated every 24 hours. **B)**. Bacterial load of double-injected flies. Mock-Primed and *Efae*-Primed flies do not differ in their bacterial load over time (Kruskal-Wallis Test: df = 6, X^2^ = 7.6661, = 0.2636). Data displays up to day 5 because of the strong survivor bias inherent to selecting flies that are still alive after that point (reference survival at day 5 and after in **Fig 1C**).

To compare *E. faecalis* priming to the priming described for *Streptococcus pneumoniae*, which was dependent on phagocytosis (Pham, et al. 2007), we performed the double injections in an *Eater* mutant background (Bretscher, et al. 2015). The hemocytes in these flies are unable to carry out bacterial phagocytosis and have cell adhesion defects in the larva, but can still mount a full Toll and IMD immune response (Kocks, et al. 2005). By comparing the *Efae*-primed to Mock-primed flies, we can observe a modest amount of immune priming, with a median survival time of 3 days and 1 day, respectively (Figure 1E). However, the *Efae-*primed flies have a shorter median survival time than the PBS/PBS controls, indicating that phagocytosis is needed to allow *Efae*-primed flies to survive as well as the double injection control.

### Priming does not increase resistance to *E. faecalis*

To measure the infection dynamics underlying both the un-primed and primed response to *E. faecalis*, we tracked bacterial load throughout the course of the infection. Infected flies were collected at 24 hour intervals after injection, homogenized, and plated in a serial dilution. As a baseline, we followed bacterial load in flies solely injected with either a high (∼30,000 CFU/fly) or low dose (∼3,000 CFU/fly) of *E. faecalis* (Figure 2A). By day 2 after injection, the bacterial loads in flies infected with a high dose were generally above 100,000 CFU/fly. This indicates that without priming, the bacterial load in flies infected with a lethal dose increases to a high plateau. In contrast, by day 1 the distribution of bacterial loads in flies initially infected with a low dose was bimodal, consistent with what has been previously reported (Duneau, et al. 2017). This suggests a subset of flies were more effectively resisting the infection and attempting to clear it, while another subset tolerated a relatively high bacterial load. The data from the low dose flies indicate two things. First, even a low dose of *E. faecalis* is not completely eliminated from the animals. Second, upon reinfection, there are likely two distinct populations of flies, harboring either a relatively high or low bacterial burden, which could alter their capability to survive a subsequent infection.

We then tested the relationship between bacterial burden and the enhanced survival seen in primed flies. Flies that are primed could increase their survival by either more efficiently clearing the infection or more effectively tolerating a chronic bacterial burden. When looking at bacterial load in double-injected flies, there was no significant difference between Mock-primed and *Efae*-primed cohorts (Kruskal-Wallis Test: p = 0.2636) (Figure 2B). Despite their significant differences in survival (Figure 1C), this does not correlate with a difference in the bacterial load between the two conditions, indicating that the improved survival of *Efae-*primed flies relative to the Mock-primed flies is due to tolerance, not resistance.

### Fat bodies show priming-specific transcription

To correlate increased survival in primed flies with transcriptional response, we measured gene expression in the fat body using RNA-seq. The fly fat body is a liver-like tissue responsible for driving an extensive transcriptional program in response to bacterial infections (DiAngelo, et al. 2009; Dionne 2014). As in previous priming setups, flies were injected twice, with samples being collected at multiple time points to assay the priming phase as well as re-infection (Figure 3A; Supplementary Table 1). To identify genes differentially expressed in response to each injection, we performed differential gene expression analysis against a non-injected, age-matched control. The response to each injection was measured after 24 hours. Genes that were differentially up-regulated only in *Efae*-primed flies were identified as “priming-specific”. As a comparison to prior work, we analyzed the expression profiles of a previously-published list of “core” immune genes in our samples and found a subset was induced upon infection in our samples (Supplementary Figure 2A) (Troha, et al. 2018).

**Figure 3.**
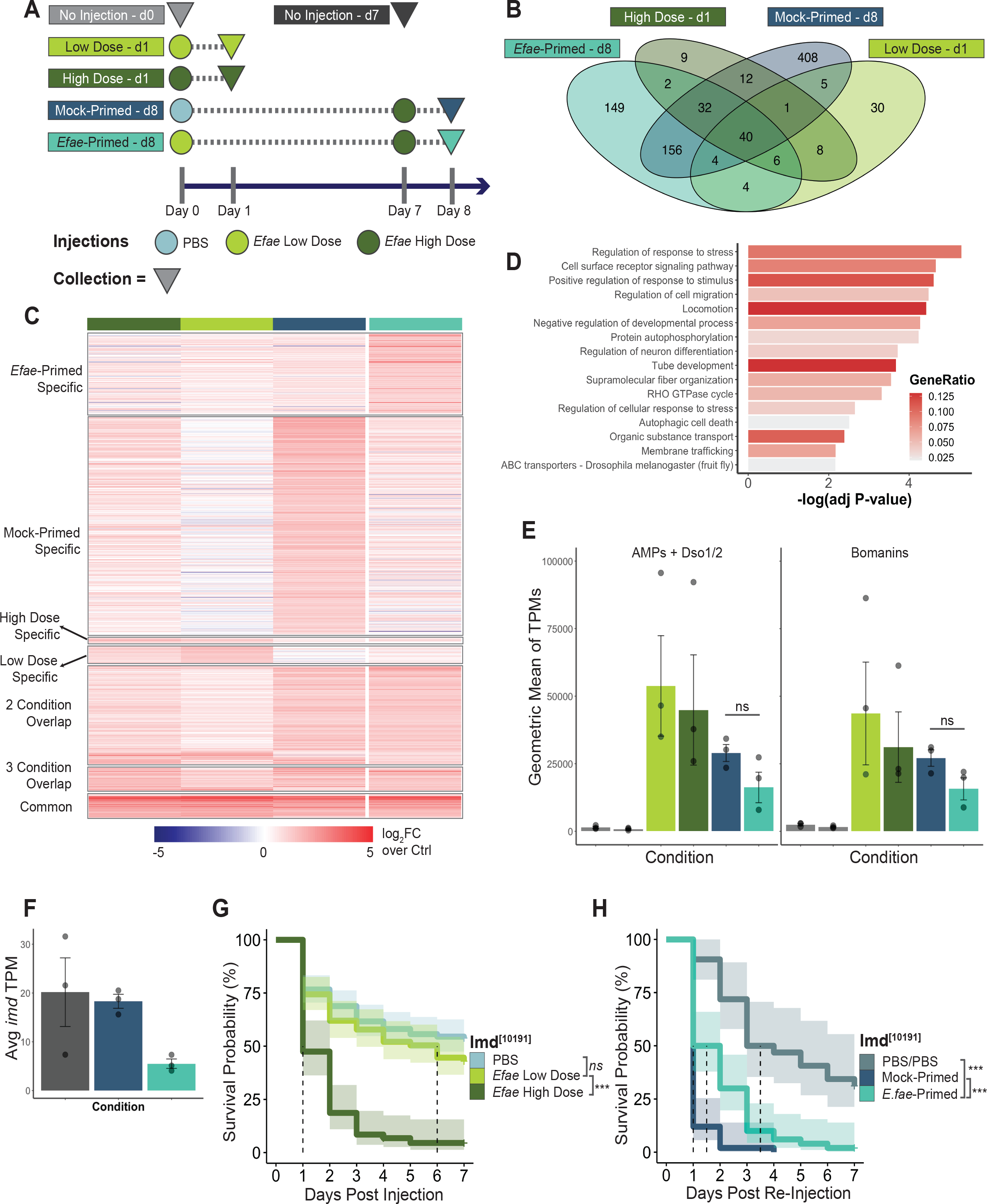
Fat bodies have a high degree of priming-specific transcriptional up-regulation. **A)**. Sample collection for RNA-seq experiments. Conditions are the same as **Figure 1A**, with the addition of age-matched, non-injected controls at Day 0 and Day 7. Circles represent injections and triangles represent time of collection. **B)**. Venn-diagram of significantly up-regulated genes (log fold change (log_2_FC) >1 & false discovery rate (FDR) <0.05) for conditions in **A** compared to age-matched controls. **C)**. Heat map of significantly up-regulated genes as corresponding to **B** (scale: log_2_FC over age-matched controls) **D)**. GO term enrichment from fat body priming-specific, up-regulated genes. **E)**. Geometric means of transcripts per million (TPMs) of core fat body *E. faecalis*-response genes across collected fat body samples. Genes are divided up by identity: [left] AMPs + Daisho 1&2 (Mock-Primed vs *Efae*-primed; Welch’s t-Test: p = 0.1835) or [right] Bomanins (Mock-Primed vs *Efae*-primed; Welch’s t-Test: p = 0.112) **F)**. Average TPMs for the gene *imd* in double-injected fat body samples. **G)**. Survival single injected *imd*-mutant flies. PBS (n = 167), *Efae* Low Dose (n = 121), and *Efae* High Dose (n = 59). PBS vs Low Dose: p = 0.098; Low Dose vs. High Dose: p < 0.0001; all survival significance testing is log rank-sum test. Dotted line represents the median survival time; shaded region indicates 95% confidence interval. **H)**. Survival of primed *imd*-mutant versus double-injected, non-primed controls (PBS/PBS: n = 55, Mock-Primed: n = 69, *Efae*-Primed: n = 42). PBS/PBS vs *Efae*-Primed: p < 0.0001; Mock-Primed vs. *Efae*-Primed: p < 0.0001.

The comparison of fat body transcription across conditions showed a high amount of *Efae* primed-specific and Mock-primed specific upregulation (149 genes & 408 genes, respectively, using an FDR cutoff of 0.05) (Figure 3B & C, full list for all conditions and overlap in Supplementary Table 2). A fraction of these genes have been previously annotated with immune functions (19 *Efae*-primed genes, ∼13%; 15 Mock-primed genes, ∼4%) (Ramirez-Corona, et al. 2021; Troha, et al. 2018). Gene ontology (GO) analysis of priming-specific up-regulation was enriched for genes related to immune response, control of response to stress, and cell surface receptor signaling (Figure 3D), consistent with the idea of bacterial sensing being essential to building a primed response (Figure 1D). Mock-primed specific GO term enrichment indicated response to stimuli, but also included genes involved specifically in response to mechanical stimuli and post-transcriptional gene regulation (Supplementary Figure 2A & Supplementary Table 2).

To delineate pathways whose component genes were upregulated in *Efae*-primed fat body versus Mock-primed fat body transcriptomes, we applied gene set enrichment analysis (GSEA) on the full transcriptome for both conditions. *Efae*-primed samples were enriched for pathways involved in protein and lipid metabolism and metabolite transport, while Mock-primed fat bodies were enriched for pathways involved in the cell cycle (Supplementary Figure 3; full analysis in Supplementary Table 3). This suggests there is metabolic reprogramming associated with priming and altered regulation of cell division in Mock-primed fat bodies. Despite the high degree of unique transcriptional activity in Mock-primed fat bodies, Mock-primed flies die more quickly than either *Efae*-primed or high dose-infected flies. This suggests that this transcriptional reaction is not necessarily advantageous for infection survival. Taken together, fat bodies showed a strong transcriptional response to infection, with a high degree of Mock-primed and *Efae*-primed-specific transcription.

We also noted that all conditions shared a set of 40 commonly up-regulated genes, which we call “core genes.” Seventeen of these core genes are known or suspected AMPs, including several *Bomanins* (*Boms*), *Daisho 1 & 2*, and the AMPs *Metchnikowin, Drosomycin, Diptericin B*, and *Baramicin A* (Supplementary Figure 2B) (Cohen, et al. 2020; Hanson, et al. 2019; Hanson, et al. 2021; Lindsay, et al. 2018). Previous experimental work has shown that survival of *E. faecalis* infection is strongly dependent on the *Bom* gene family (Clemmons, et al. 2015). Flies lacking 10 out of the 12 *Boms* succumb to a single *E. faecalis* infection as quickly as flies that lack Toll signaling. Bacterial load data indicates that flies lacking either these 10 *Boms* resist an individual *E. faecalis* infection more weakly than wild type flies. Conversely, flies with deletions of several AMPs (4 Attacins, 2 Diptericins, Drosocin, Drosomycin, Metchnikowin, and Defensin) or *Baramicin A* show only modest decreases in survival of *E. faecalis* infections (Hanson, et al. 2019; Hanson, et al. 2021).

Given their differing effects on *E. faecalis* infection survival, we decided to analyze the expression patterns of the core *Boms* separately from the other core known or suspected AMPs. We summarized the expression patterns of each gene group using a geometric mean of transcripts per million (TPMs). When comparing the geometric means of the core *Boms*, we found no significant difference in expression between the Mock-primed and *Efae*-primed flies (Welch t-test: p = 0.112) (Figure 3E, left). Likewise, a comparison of the geometric means of expression levels for the core AMP or AMP-like genes yielded no significant difference between the Mock-primed and *Efae*-primed flies (Welch t-test: p = 0.184) (Figure 3E, right). This indicates that primed fat bodies are not necessarily increasing the amount of transcripts associated with bacterial resistance, consistent with the lack of increased bacterial clearance for *Efae*-primed relative to Mock-primed flies in Figure 2B.

### Loss of IMD negatively impacts the fly’s ability to prime against *E. faecalis*

We also observed priming-specific down-regulation of *imd* (Figure 3F), which led us to consider the role of IMD signaling in the priming response. While IMD signaling is canonically associated with response to Gram-negative bacterial infections, it is also connected to regulation of the MAPK-mediated reactive oxygen species production and wound response, as well as a generalized stress response (Ragab, et al. 2011; Myllmäki, et al. 2014). We first hypothesized that the downregulation of *imd* in *Efae-*primed flies might lead to lower expression levels of IMD-responsive AMPs, perhaps as a way to avoid transcribing genes that do not contribute to the animal’s survival of the Gram-positive *E. faecalis* infections. However, the IMD-responsive AMPs were not down-regulated in a priming-specific manner (Supplementary Figure 2C & D).

To further explore the role IMD signaling plays in a primed immune response, we tested survival of an *imd* mutant (Pham, et al. 2007) to single and double injections (Figure 3 G & H, Supplementary Figure 2E & F). As has been previously shown, the *imd* mutant showed a dose-dependent response to *E. faecalis* infection with similar survival to a single PBS injection and a low dose of *E. faecalis* (Figure 3G), indicating that loss of the pathway did not impact the ability of the fly to respond to an *E. faecalis* infection. However, when subjecting the flies to dual injections, we observed a significant, though not total, loss of priming ability in these *imd*-mutant flies (Figure 3H). *Efae*-primed flies still survive a second injection more effectively than Mock-primed flies, but less successfully than control flies twice injected with sterile PBS. Together, this demonstrates that while the loss of the IMD pathway does not impact the survival of the flies with a single bacterial infection, it does negatively impact survival in animals that have been infected more than once. This suggests that there are distinct differences in use of signaling pathways between animals with one versus two infections.

### Hemocytes act as potential signal relayers in a primed immune response

Using the same approach as in fat bodies, we determined priming-specific transcription in adult hemocytes (Supplementary Figure 4A, full list of up-regulated and down-regulated genes in Supplementary Table 4). Hemocytes have several roles in the immune response, including bacterial phagocytosis, pathogen sensing, and signaling. Compared to fat bodies (Figure 3B), hemocytes showed a low amount of priming-specific up-regulation, with only 17 genes specifically up-regulated in the *Efae*-primed condition (Figure 4A, Supplementary Figure 4B). There were also 458 genes specifically up-regulated in *Efae* High hemocytes, indicating that the hemocyte transcriptional response to *E. faecalis* infection depends on the dose, previous injection state, and age of the animal. A GO term analysis reveals that many of these high dose specific genes are involved in immune response, as expected, and also regulation of metabolic processes (Supplementary Figure 4C). This Most of these genes are poorly characterized or functionally unrelated (Supplementary Table 4). analysis indicates that, in contrast to the fat body, hemocytes only upregulate a small number of genes in the primed condition.

**Figure 4.**
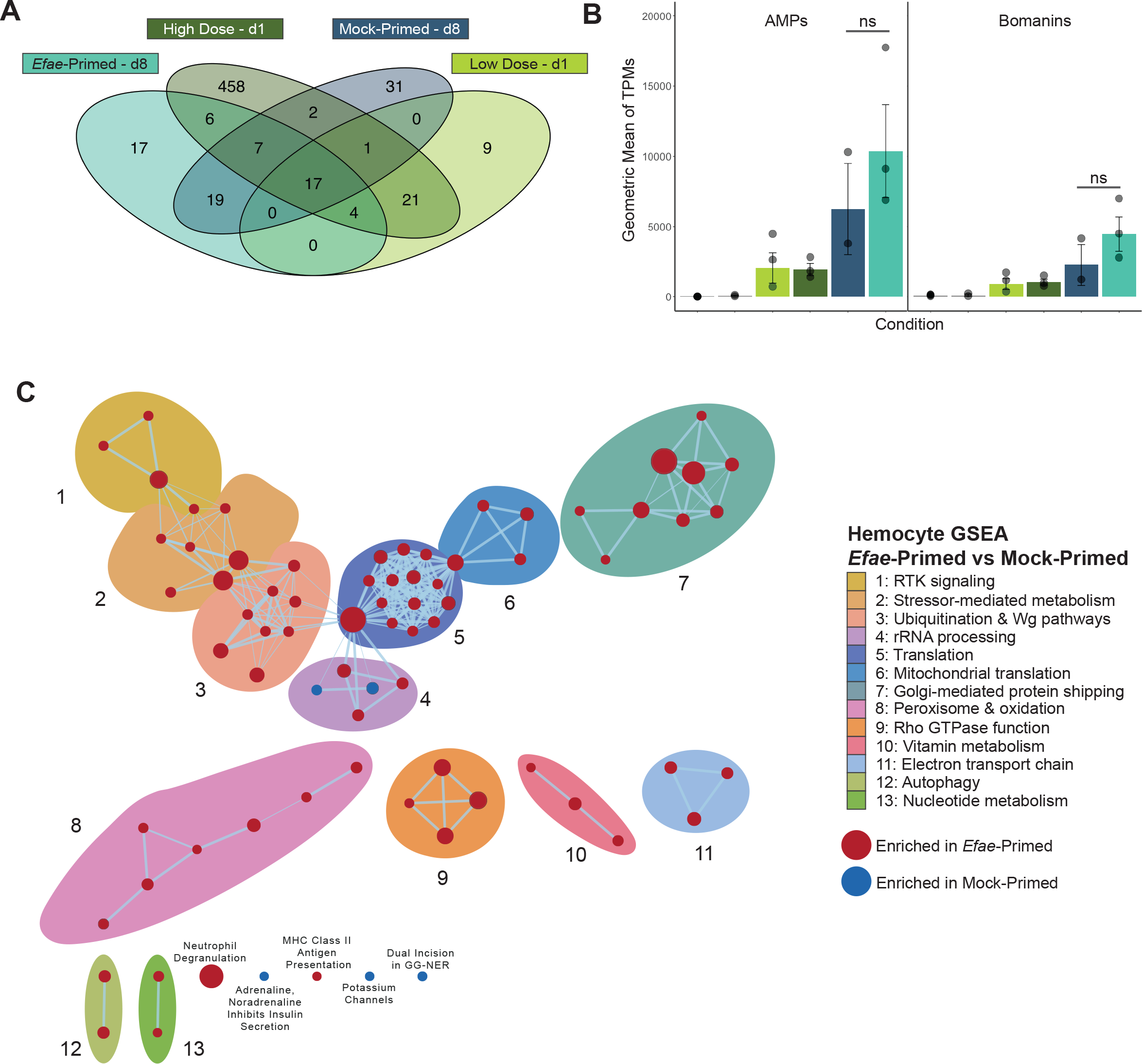
Hemocytes do not significantly increase effector expression when primed, but differentially activate metabolic pathways. **A)**. Venn diagram of significantly up-regulated (log_2_FC >1 & FDR <0.05) genes for hemocytes collected at the same conditions as **Fig 3A. B)**. Geometric means of TPMs of core hemocyte *E. faecalis*-response genes across collected hemocyte samples. Genes are divided up by identity: [left] AMPs (Mock-primed vs *Efae*-primed; Welch’s t-Test: p = 0.4391) or [right] Bomanins (Mock-primed vs *Efae*-primed; Welch’s t-Test: p = 0.3773). **C)**. Gene set enrichment analysis for *Efae*-Primed versus Mock-Primed hemocytes. This visualization represents relationships between statistically significant terms (FDR < 0.05), manually curated with clusters that summarize the relationships between terms. Full results are found in **Supplementary Table 5**.

Of the 17 core genes up-regulated in all conditions in hemocytes, 11 of them (∼64%) overlapped with the 40 core genes found in fat bodies (Supplementary Figure 4D & Supplementary Table 4). These hemocyte core genes were identified to be the overlapping up-regulated genes between all four conditions that assayed immune response 24 hours after either single or double injection. Among these were several Bomanins, *Drosomycin, SPH93, IBIN*, and *Metchnikowin-like*, implying a role for these genes in response to *E. faecalis* infection in both hemocytes and fat body. As with our fat body data, we again separately analyzed the levels of expression of the AMPs versus bomanin effectors for hemocytes. When comparing the geometric means of the expression levels of the core *Boms*, we found no significant difference in expression between the Mock-primed and *Efae*-primed flies (Welch t-test: p = 0.3773) (Figure 3B, right). Likewise, a comparison of the geometric means of expression levels for the core AMP genes yielded no significant difference between the Mock-primed and *Efae*-primed flies (Welch’s t-test: p = 0.4391) (Figure 3B, left). This indicates that, similar to the comparison between *Efae-*primed and Mock-primed fat bodies, transcripts associated with bacterial resistance are not specifically up-regulated in primed hemocytes.

Given the diverse functions of hemocytes in immune response, we decided to use GSEA to systematically delineate priming-enriched pathways (Figure 3C, full GSEA analysis in Supplementary Table 5). This analysis of hemocyte transcription in *Efae*-primed samples versus Mock-primed samples indicated a wider picture of metabolic reprogramming (Clusters 2, 6, 8, 10, 11, and 13) and altered protein production (Clusters 4, 5, 6, and 7) in the primed samples. Though not clustered with other terms, there was also enrichment for genes involved in antigen-presenting functions in mammalian orthologs. To more fully understand the role hemocytes could be playing in modulating a primed response, we synthesize several of our observations. The decreased priming ability in *Eater* mutants indicates that bacterial phagocytosis is necessary for immune priming (Figure 1F), but we do not find an increase in bacterial clearance in primed re-infection (Figure 2B). Consistent with this observation, we also do not see elevated transcription of either the *Boms* or other known or suspected AMPs typically associated with bacterial clearance (Figure 4B). Transcriptional profiling of the hemocytes point to changes in regulation of metabolism and protein production (Figure 4C) that may also contribute to the enhanced survival of primed animals. Together these observations suggest that, in the primed condition, the primary role of bacterial phagocytosis is to initiate bacterial sensing and subsequent signal transduction (Nehme, et al. 2011; Gold & Brückner 2014).

### Several Toll effectors loiter into re-infection, but Toll signaling is not needed for immune priming

We further leveraged our transcriptomic data to identify genes that loiter from the first infection into reinfection (Figure 5A). We defined loitering genes as those that wereup-regulated both 1 day and 6 days after a low dose infection (*Efae* Low-d1 & *Efae* Low-d7) and 1 day after the subsequent high dose infection (*Efae*-primed-d8). Fat bodies had 14 genes that were identified as loitering (Figure 5B), while hemocytes only had two (Figure 5C). For fat bodies, 13 of the 14 (∼93%) loitering genes overlapped with the identified core *E. faecalis* response genes (Figures 3B & C; annotated in Supplementary Table 2). Most of these genes are either known or suspected AMPs, and the list also includes a recently-characterized lncRNA (lncRNA:CR33942) that can enhance the Toll immune response (Zhou, et al. 2022). The fat body loitering genes are largely Toll-regulated.

**Figure 5.**
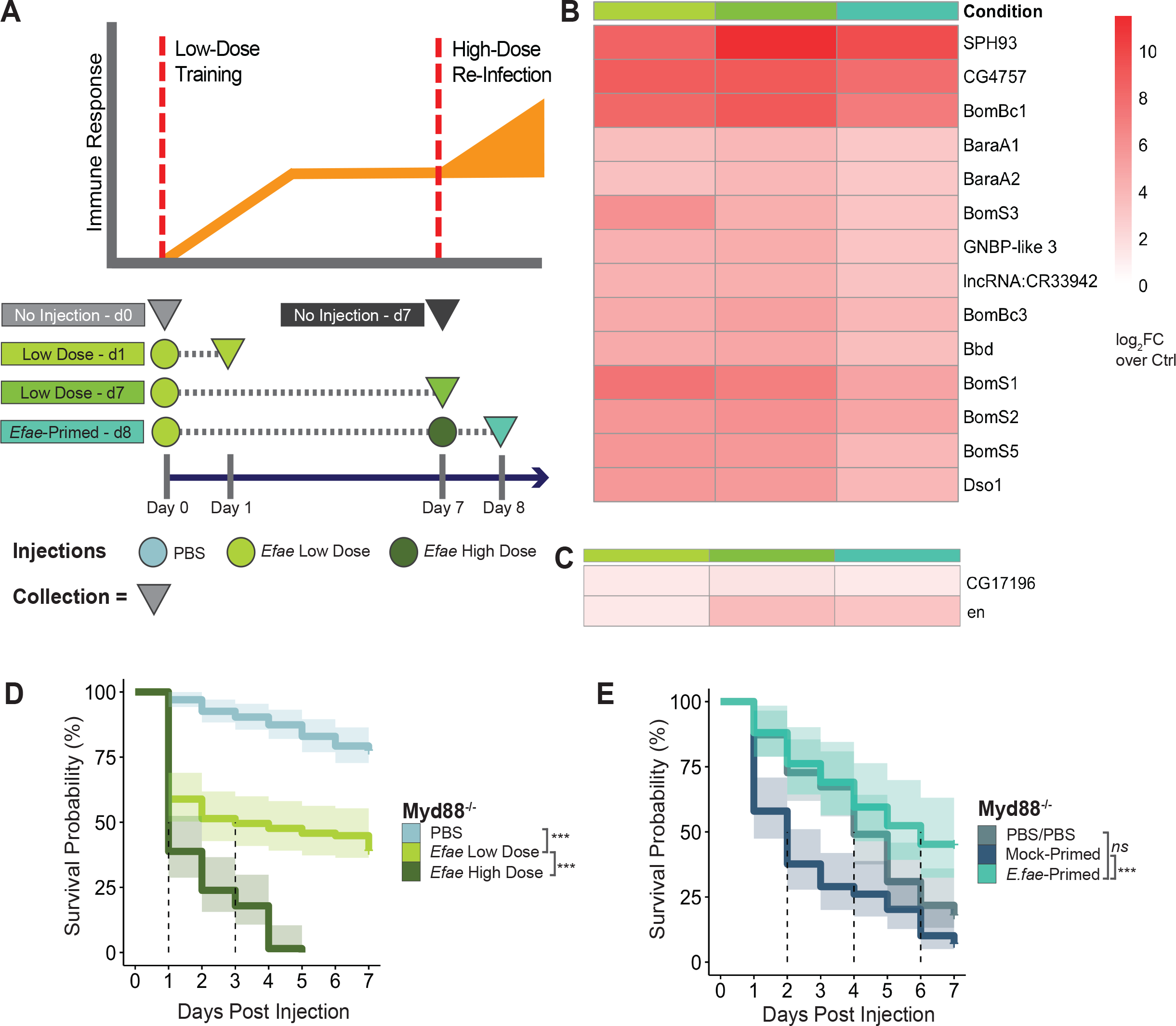
Toll effector genes loiter throughout *E. faecalis* immune priming. **A)**. Schematic of immune loitering from priming into re-infection. Experimental conditions are the same as **Figure 1A**, with the addition of age-matched, non-injected controls at Day 0 and Day 7 as well as an additional time point at Day 7 for collection of samples late in priming. Circles represent injections and triangles represent time of collection **B)**. Immune loitering genes in fat bodies (scale: log_2_FC over age-matched controls). **C)**. Immune loitering genes in adult hemocytes (scale: log_2_FC over age-matched controls). **D)**. Survival of single injected *Myd88*-mutant flies. PBS (n = 135), *Efae* Low Dose (n = 107), and *Efae* High Dose (n = 67). PBS vs Low Dose: p < 0.0001; Low Dose vs. High Dose: p < 0.0001; all survival significance testing is log rank-sum test. **E)**. Survival of primed *Myd88*-mutant versus double-injected, non-primed controls (PBS/PBS: n = 55, mMock-Primed: n = 69, *Efae*-Primed: n = 42). PBS/PBS vs *Efae*-Primed: p = 0.021; Mock-Primed vs. *Efae*-Primed: p < 0.0001.

To further investigate the role Toll signaling is playing in creating a primed response to *E. faecalis*, we assayed infection response in flies with a *Myd88* mutation that eliminates Toll signaling (Figure 5D) (Charatsi, et al. 2003). In the single injection conditions, we continued to see a dose-dependent effect on survival, with expected increased lethality when compared to our immune-competent control (Supplementary Figure 5A) (Clemmons, et al. 2015; Hanson, et al. 2019). When assaying for survival against double-injected conditions, we found that *Myd88* mutants were still able to effectively prime against *E. faecalis* re-infection (Figure 5E). Despite lacking canonical Toll-mediated immune signaling, these mutants were able to respond to double-injections and mount a primed immune response, with equivalent survival between the *Efae*-primed flies and the control flies injected twice with PBS (Supplementary Figure 5B). This indicates that immune priming against the Gram-positive *E. faecalis* does not strictly require Toll signaling.

### Potentiated recall gene expression plays a minor role in *E. faecalis* immune priming

In addition to priming-specific and loitering genes, we were also identified “recall response genes” (Melillo et al. 2018). These genes were defined as genes that are up-regulated in response to an initial low dose infection, turned off 6 days later, and up-regulated more strongly in response to a subsequent infection (Figure 6A). In fat bodies, we identified 7 recall genes (Figure 6B), and we did not identify any recall genes in hemocytes. Of these few fat body recall genes, we found two Polycomb interacting elements (*jing* & *cg*) and a component of the Mediator complex (*MED23*), suggesting a potential role for transcriptional regulation. However, we did not find a strong role for recall transcription in our experiments.

**Figure 6.**
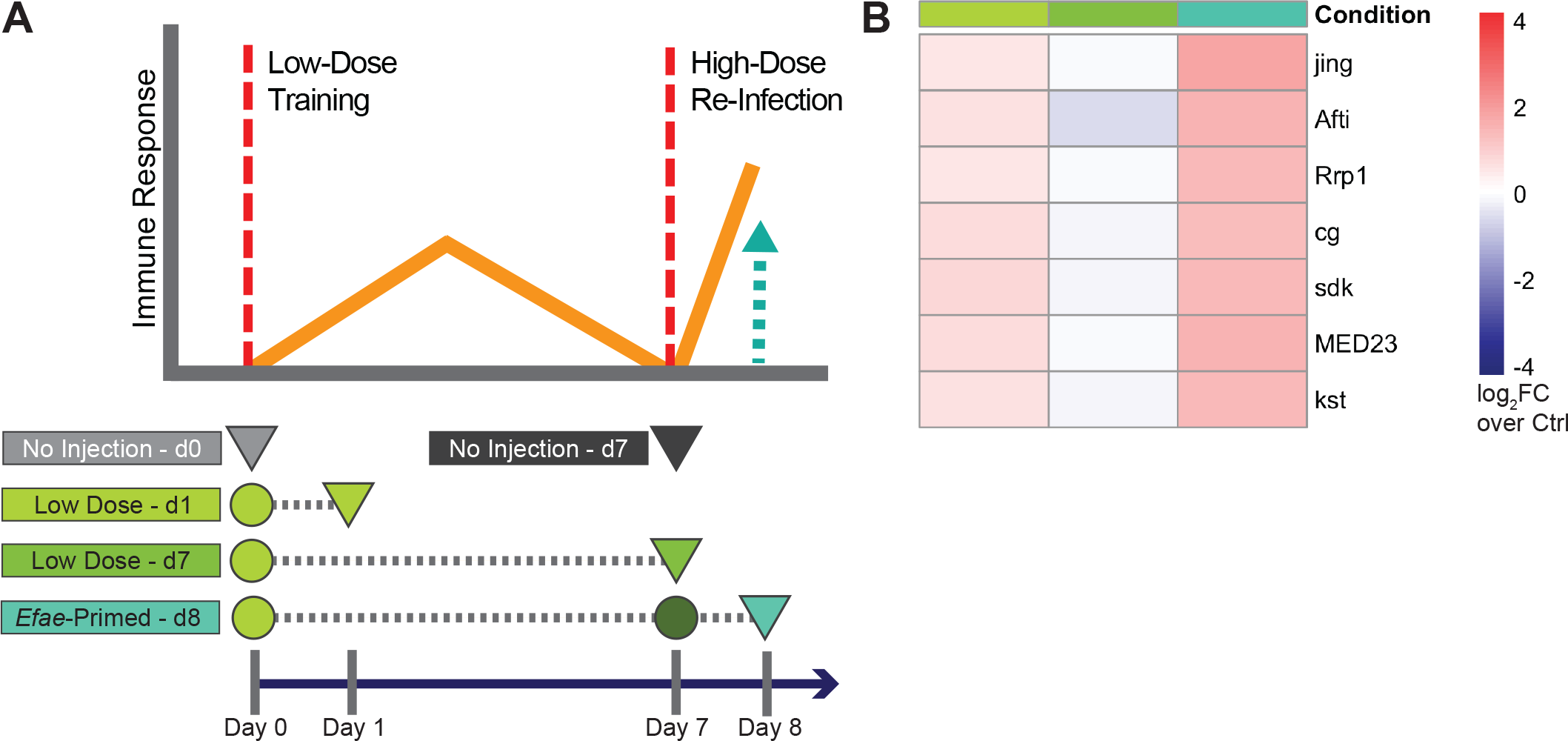
Few potentiated genes are recalled in *E. faecalis* immune priming. **A)**. Schematic of immune recall response. **B)**. Potentiated recall genes in fat bodies (scale: log_2_FC over age-matched controls).

## Discussion

In this study we have shown the transcriptional underpinnings of a primed immune response against *Enterococcus faecalis* infection in *Drosophila melanogaster*. We demonstrated that a low dose of *E. faecalis* can prime the flies to better survive a high dose infection at least 7 days later, and the increase in survival is not linked to more effective clearance of the bacteria. When comparing *Efae*-primed and Mock-primed animals, we found that the transcriptional profiles of antimicrobial peptides and Bomanins do not differ between the two conditions in either the fat body nor the hemocytes. However, there are ample transcriptional differences between the conditions, and GSEA analysis points to differences in cell cycle regulation and metabolic response. When testing priming ability in *imd* and *Myd88* mutants, we found that these mutants have unexpected effects in the double injection conditions – *imd* mutants prime less effectively than wild type flies, while *Myd88* mutants show no apparent loss of priming ability.

There are previous studies of immune priming in flies, which taken together with this work paint a more complete picture of the phenomenon. One of the early descriptions of immune priming in *D. melanogaster* found a phagocytosis-dependent, AMP-independent priming response against *Streptococcus pneumoniae* (Pham, et al. 2007). Our study uses a different Gram-positive microbe, but a similar re-infection timescale. Similar to that study, we find that phagocytosis is needed to mount a primed immune response, as was demonstrated by the impaired priming in the Eater mutant flies. We also corroborated that survival is not correlated with AMP production. However, Pham et al. found that primed flies resist *S. pneumoniae* more effectively than naive flies, while our *Efae*-primed flies appeared to rely on immune tolerance to enhance survival. It is possible that this difference is due to the increased virulence of the pathogen, *S. pneumoniae*, which can kill a wild type fly with a relatively low dose of 3,000 CFU, relative to *E. faecalis*. The difference could also be due to the specificity of the host’s primed response to different pathogens. In sum, these findings suggest that there may be multiple, bacteria-specific priming mechanisms.

Another study found that sterile wounding 2 days, but not 7 days, prior to infection with *E. faecalis* conferred some level of ROS-mediated protection (Chakrabarti, et al. 2020). This study’s assay most closely matches our Mock-primed re-infections, and we also did not see enhanced survival when the wounding occured 7 days prior to the infection. This indicates that the protection conferred from sterile wounding is effective in the short-term (i.e. 2 days), but not in the long-term (i.e. 7 days). However, both this study and our observations support the idea that hemocytes activate new functions in response to prior stimuli exposure (as was found in Weaver, et al. 2016, as well). Finally, a study looking at the effects of chronic bacterial infection did not find immune priming with *E. faecalis* when using the same re-injection time points (Chambers, et al. 2019). However, in that study flies were injected with two low-doses (∼3,000 CFU/fly) and injected first in the abdomen and second in the thorax. This suggests a dose-dependent and/or injection site-dependent effect on priming ability.

One of the most surprising findings of this study is the priming responses found in the *imd* and *Myd88* mutant flies. As others have previously reported, our work demonstrates that the elimination of the IMD pathway does not affect the fly’s survival against a single low dose infection of *E. faecalis*, while the elimination of Toll signaling greatly reduces the fly’s survival of the same infection. This is consistent with the well-described sensing of Gram-positive bacteria via Toll signaling and Gram-negative bacteria via IMD signaling (Buchon, et al. 2014). However, we find that *imd* mutants lose some, though not all, of their priming capacity, while *Myd88* mutants have similar survival between flies injected twice with PBS or *Efae*-primed flies. The requirement of *imd* for survival was surprising for two reasons: first because IMD signaling has not been implicated in the survival of Gram-positive bacteria (or priming, in the case of *S. pneumoniae* in Pham, et al. 2007), and second, because we saw down regulation of the *imd* gene in the fat body primed transcriptome. This suggests while downregulation of *imd* may be useful in priming, complete eradication of the pathway in the animal removes some priming ability. This could be due to the role the IMD pathway plays in modulating other key immune response pathways such as JAK/STAT, JNK, and MAPK signaling (Kleino & Silverman 2014).

We were also surprised to see the dispensability of Toll signaling for priming. Toll signaling plays a key role in surviving Gram-positive infections, and virtually all of the loitering genes we found here are known Toll targets. One possible explanation of this observation is that *Myd88* mutants show markedly lower survival of the initial low dose *E. faecalis* infection. This implies that, when we select survivors to re-infect 7 days later, this may be representative of a specific subset of flies with an advantage that allows them to survive the initial infection despite the lack of a Toll response.

While our data did not indicate a difference in bacterial clearance between *Efae*-primed and Mock-primed flies (Figure 2B), we acknowledge the possibility that the number of bacteria remaining in the animal from the initial infection may affect priming responses. As has been previously noted (Duneau, et al. 2017), we found variability in the bacterial burden during the initial low dose infection, consistent with some flies more effectively resisting infection than others (Figure 2A). Chronic infections tend to lead to low-level activation of the immune response throughout the animal’s lifetime, causing expression of immune effectors that can loiter into re-infection and and may contribute to enhanced survival (Chambers, et al. 2019). It is not yet clear what effect the intensity of a chronic infection would have on an priming ability, but it should be considered in the future. It is possible that a more severe chronic infection could either put the animal in a heightened state of “readiness” for a new infection or exhaust its resources.

Our data implies a major role for metabolic reprogramming in mediating a primed immune response against *E. faecalis*. Given the high energetic cost of mounting an immune response, it is logical to imagine immune priming as a more efficient re-allocation of metabolic resources to fine tune an immune defense strategy in a short-lived animal (as discussed in Lazarro & Tate 2022; Schlamp, et al. 2021). Interestingly, evidence of metabolic shifts was not just relegated to the fat body (Supplementary Figure 3), which acts as the site of integration for metabolic and hormonal control, but was found to be the case with hemocytes, as well (Figure 4C). Similarly, in mammalian trained immunity where metabolic reprogramming drives epigenetic changes in innate immune cell chromatin(Fanucchi, et al. 2021). Further characterization of *Drosophila* immune priming could explore the extent of differential metabolite usage when mounting a primed immune response and whether the transcriptional differences observed are encoded through epigenetic reprogramming of histone mark deposition, akin to what is observed in mammalian systems. Our study lays the groundwork for understanding the interplay between a physiological primed immune response and the transcriptional regulatory logic defining it.

## Methods

### Fly Strains

Experiments, unless otherwise indicated, were performed using 4 day old Oregon-R male flies. Eater mutants are described in Bretscher et al. (2015) and were obtained from the Bloomington Stock Center (RRID:BDSC_68388). These flies knocked out the *eater* gene through homologous recombination that replaced 745bp of the TSS, exons 1 and 2, and part of exon 3 with a 7.9 kb cassette carrying a w[+] gene. Imd1091 flies were provided by Neal Silverman. They were generated by creating a 26bp deletion at amino acid 179 that creates a frameshift mutation at the beginning of the death domain in *imd* (Pham 2007). Myd88[kra-1] flies were provided by Steve Wasserman and Lianne Cohen. This line was created by excising 2257bp of the *Myd88* gene spanning the majority of the first exon and inserting a P-element (Charatsi 2003). Stable lines were balanced against a CyO balancer with homozygous mutant males being selected for injections. Flies were housed at 25°C with standard humidity and 12 hr-light/12 hr-dark light cycling.

### Injections

All bacterial infections were done using a strain of *Enterococcus faecalis* originally isolated from wild-caught *Drosophila melanogaster* (Lazarro 2006). Single colony innoculumns of *E. faecalis* were grown overnight in 2mL BHI shaking at 37°C. 100uL of overnight *E. faecalis* innoculumn was then added to 2mL fresh BHI and grown shaking at 37°C for 2.5 hours before injections in order to ensure it would be in the log-phase of growth. Bacteria was then pelleted at 5,000 rcf for 5 minutes, washed with PBS, re-suspended in 200uL PBS, and measured for its OD600 on a Nanodrop. Flies were injected with either PBS, *E. faecalis* at OD 0.05 for low dose experiments (∼3,000 CFU/fly), or *E. faecalis* at OD 0.5 for high dose experiments (∼30,000 CFU/fly). Due to the high heat resistance of *E. faecalis*, heat-killed inoculums were produced by autoclaving 10mL cultures that were in log-phase growth. Successful heat-killing was determined by streaking 50uL on a BHI plate and checking it had no growth. Adult flies were injected abdominally using one of two high-speed pneumatic microinjectors (Tritech Research Cat. # MINJ-FLY or Narishige IM 300) with a droplet volume of ∼50nL for both PBS and bacterial injections. Injections into a drop of oil on a Lovins field finder were used to calibrate the droplet volume. Injections were performed in the early afternoons to control for circadian effects on immune response. Flies were not left on the CO_2_ pad for more than 10 minutes at a time. Injected flies were housed in vials containing a maximum of 23 flies at 25°C with standard humidity and 12 hr-light/12 hr-dark light cycling.

### Survival Assays

To track survival, flies were observed every 24 hours at the time they were injected. Media was changed every three days with flies being exposed to CO_2_ for no more than two minutes between vial transfers. Survival was modeled and analyzed using a log rank-sum test and visualized using the R packages survival and surminer.

### Dilution Plating

Single flies were suspended in 250uL PBS and homogenized using an electric pestle. The homogenate was then serially diluted five-fold and plated on BHI plates and left to grow in aerobic conditions for two days at 25°C. Using this method there was little to no background growth of the natural fly microbiome. Images were then taken of each plate using an iPhone XR and analyzed using ImageJ with custom Python scripts to calculate colony forming units (CFU) per fly. Plotting was done using the R package ggplot2 (Wickham 2016).

### Hemocyte Isolation

For each biological replicate, 20 flies were placed in a Zymo-Spin P1 column with the filter and silica removed along with a tube’s-worth of Zymo ZR BashingBeads. Samples were centrifuged at 10,000 rcf at 4°C for one minute directly into a 1.5mL microcentrifuge tube containing 350uL TriZol (Life Technologies) (schematic in Supplementary Figure 4A). Samples were then snap frozen and stored at -80°C for future RNA extraction.

### Fat Body Isolation

Each biological replicate consisted of 3 extracted fat bodies. Flies were anesthetized with CO_2_ and pinned with a dissection needle at the thorax, ventral side up, to a dissection pad. The head, wings, and legs were then removed using forceps. Using a dissection needle, the abdomen was carefully opened longitudinally and the viscera removed using forceps. The remaining abdominal filet with attached fat body cells was then removed from the thorax and transferred to a 1.5mL microcentrifuge tube on ice containing 350uL TriZol. Samples were then snap frozen and stored at -80°C for future RNA extraction. Dissection of fat bodies includes some level of testes and sperm contamination, which was monitored by tracking expression of sperm-related genes in RNA-seq libraries and throwing out any libraries that have relatively high expression of said genes (Supplementary Figure 6).

### RNA-seq Library Preparation

RNA from either fat bodies or hemocytes was extracted using a Zymo Direct-zol RNA Extraction kit and eluted in 20uL water. Libraries were prepared using a modified version of the Illumina Smart-seq 2 protocol as previously described (Ramirez-Corona 2021). Libraries were sequenced on an Illumina Next-seq platform using a NextSeq 500/550 504 High Output v2.5 kit to obtain 43bp paired-end libraries.

### Differential Gene Expression Analysis

Sequenced libraries were quality checked using FastQC and aligned to *Drosophila* reference genome dm6 using Bowtie 2 (Langmead & Salzberg 2012). Counts were generated using the subread function featureCounts. Counts were then loaded into EdgeR (Robinson

2010), libraries were TMM normalized, and genes with CPM < 1 were filtered out. Full code used in downstream analysis can be found at https://github.com/WunderlichLab/ImmunePriming-RNAseq.

Priming-Specific Transcription Analysis

To identify priming-specific up-regulation, we first identified genes that were significantly up-regulated (log_2_FC>1 & FDR<0.05) in each condition that assayed for immune response 24 hours after infection (i.e. *Efae* Hi Dose-d1, *Efae* Low Dose-d1, *Efae* Mock-primed-d8, and *Efae*-primed-d8) (the effect of modulating significance and log_2_FC cut-offs can be seen in Supplementary Figure 7). These gene lists were then compared to each other for overlap.

Genes that were only up-regulated in *Efae*-primed-d8 samples, but in no other condition were labeled as “priming-specific”. Average expression of AMPs and *Bomanins* was calculated by taking the geometric mean of TPMs of the respective gene lists. In this way we could account for the effects highly-expressed genes would have on skewing the overall average. Significant differences between conditions were calculated using a Welch’s t-test.

### Immune Loitering Analysis

To determine genes that were continuously being expressed throughout initial immune priming into re-infection, we focused on the transcription in samples assayed at *Efae* Low-d1, *Efae* Low-d7, and *Efae*-primed-d8. We first selected genes that were expressed at the above time points relative to a non-stimulated, age-matched control (log_2_FC >0). We then filtered that shortlist on the following conditions: genes had to significantly up-regulated at *Efae* Low-d1 compared to its age-matched control (log_2_FC>0 & FDR<0.05), genes had to significantly up-regulate at *Efae*-Primed-d8 compared to its age-matched control (log_2_FC>0 & FDR<0.05), and genes had to either stay at similarly expressed levels or increase in expression between *Efae* Low-d7 and *Efae*-primed-d8 compared to their age-matched controls (log_2_FC≥0).

### Potentiated Recall Response Analysis

We termed genes as being “recalled” if they were initially transcribed during priming (*Efae* Lo-d1 log_2_FC over age-matched control > 0.5), ceased being expressed by the end of priming (*Efae* Lo-d7 log_2_FC over age-matched control ≤ 0), and were then re-expressed upon re-infection (*Efae*-primed-d8 log_2_FC over age-matched control > 0.5 & FDR < 0.1). Our significance threshold had to be somewhat relaxed for expression after re-infection in order to detect any recalled gene expression at all. To delineate genes that were truly re-activating transcription in a potentiated manner (i.e. at a higher level upon re-infection as compared to when they were initially expressed during priming), we also filtered on the conditional that log_2_FC over age-matched controls had to be higher in *Efae*-primed-d8 versus *Efae* Low-d1. Finally, to identify genes that were recalled only in our primed samples, we further filtered on the condition that genes had to have a log_2_FC ≤ 0 over age-matched controls for Mock-primed-d8 samples.

### GO Term Enrichment

All GO Term Enrichment was done using Metascape’s online tool (Zhou 2019) and plotted using custom ggplot2 scripts.

### Gene Set Enrichment Analysis

Gene set enrichment analysis was run using the GSEA software v. 4.2.3 (Subramanian 2005). *Drosophila*-specific gene matrices for both KEGG and Reactome-based GSEA alayses were taken from Cheng 2021. TMM-normalized TPMs were extracted from EdgeR analysis and used as input for two-condition comparisons using GSEA software. Due to the low number of replicates (< 7 replicates per condition), analysis was run using a gene set permutation. Full tabular results are found in Supplementary Tables 3 & 5. Analysis results were then visualized using Cytoscape (Node Cutoff = 0.1 FDR; Edge Cutoff = 0.5) and clusters describing the mapping manually curated.

## Acknowledgements

We would like to thank S. Wasserman, N. Silverman, and Bloomington Stock Center for fly strains; B. Lazarro for bacterial strains; A. Mortazavi, C.J. McGill, and H.Y. Liang for access to their sequencing core and technical assistance with library preparation. We would like to thank L. Cohen and B. Ramirez-Corona for constructive discussion on this work. This work was funded by NSF grant MCB-1953312/2223888 (to Z.W.) K.C. is an NIH-IMSD Fellow and an NSF-GRFP Fellow.

## Author Contributions

Z.W. and K.C. conceptualized and designed the experiments. K.C. did the survival experiments, injections, RNA-seq experiments, and analyzed the data. D.S.H. did the bacterial load experiments and helped analyze that data. D.M. did the heat-killed *E. faecalis* experiments. K.C. and Z.W. wrote the manuscript.

## Supplementary Table Legends

**Supplementary Table 1:** Sequencing information for fat body and hemocyte RNA-seq

**Supplementary Table 2:** Lists of up-regulated genes specific to each fat body condition assayed in Figure 3, common between all fat body conditions, and specifically down-regulated in *Efae*-primed-d8 fat bodies.

**Supplementary Table 3:** Gene set enrichment analysis for *Efae*-primed vs Mock-primed fat bodies. Clustering and terms are shown in Supplementary Figure 3. This represents the tabular output directly from the GSEA software v. 4.2.3 (Subramanian 2005).

**Supplementary Table 4:** Lists of up-regulated genes specific to each hemocyte condition assayed in Figure 4, common between all hemocyte conditions, specifically down-regulated in *Efae*-primed-d8 fat bodies, and overlap between common *Efae*-response genes in fat bodies and hemocytes.

**Supplementary Table 5:** Gene set enrichment analysis for *Efae*-primed vs Mock-primed hemocytes. Clustering and terms are shown in Figure 4C. This represents the tabular output directly from the GSEA software v. 4.2.3 (Subramanian 2005).

