## Supplementary Figure 1 for "Drosophila immune priming to Enterococcus faecalis relies on immune tolerance rather than resistance"

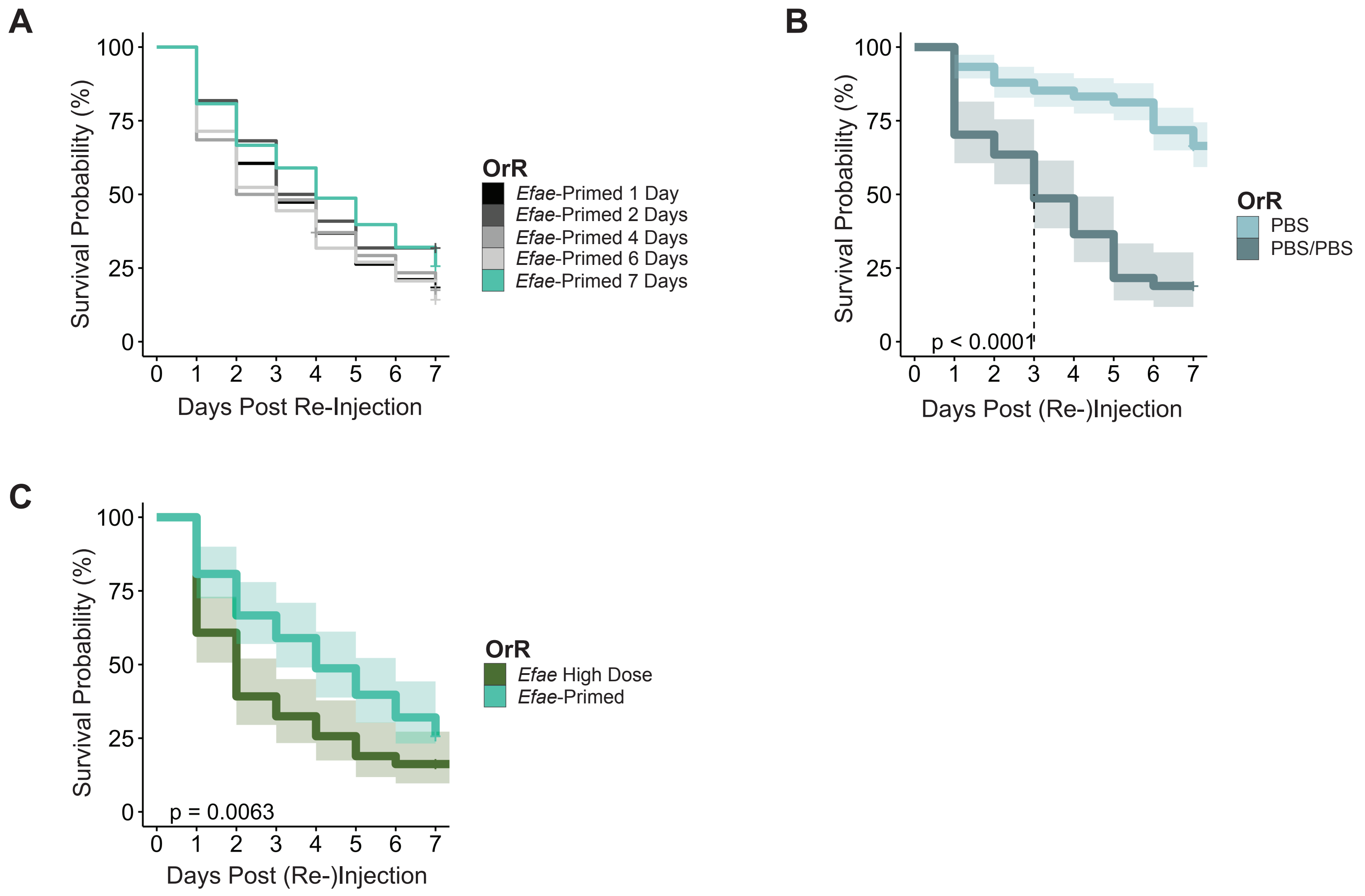


**Supplementary Figure 1: Dynamics of *E. faecalis* priming and double-injection survival in OrR flies**

**A).** Survival is similar in flies allowed to prime with a low-dose of *E. faecalis* (~3,000 CFU/fly) for varying amounts of time before re-infection with a high dose of *E. faecalis* (~30,000 CFU/fly) (n: 1 Day = 38, 2 Days = 22, 4 Days = 54, 6 Days = 63, 7 Days = 78). **B).** There is a aignificant difference in survival (log-rank sum test, $p<0.0001$) in OrR flies injected once with PBS (PBS, n = 149) or twice with PBS with seven days of rest between repeated injections (PBS/PBS, n = 74). Dotted lines indicate median survival time; shaded regions indicate 95% confidence intervals. **C).** There is a significant difference in survival (log-rank sum test, $p=0.0063$) in OrR flies injected once with a high dose of *E. faecalis* (*Efae* High, ~30,000 CFU/fly, n = 74) versus primed with a low dose of *E. faecalis* for seven days and then re-infected with a high dose of *E. faecalis* (*Efae*-Primed, n = 78). Data are the same as **Figure 1B-C**, replotted for comparison.
