## Supplementary Figure 2 for "Drosophila immune priming to Enterococcus faecalis relies on immune tolerance rather than resistance"

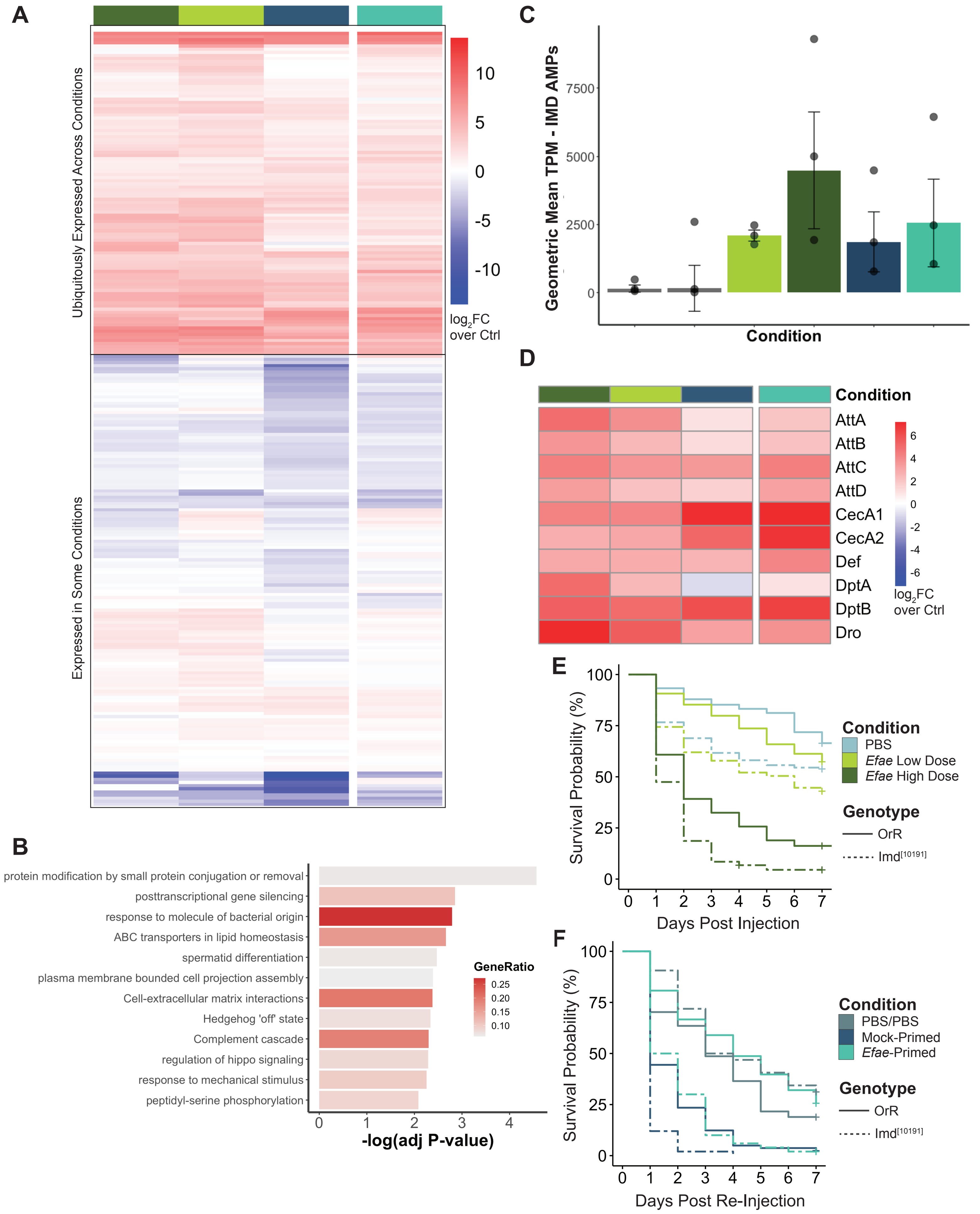
A
Ubiquitously Expressed Across Conditions
Expressed in Some Conditions
10
5
0
-5
-10
log2FC over Ctrl
B
protein modification by small protein conjugation or removal
posttranscriptional gene silencing
response to molecule of bacterial origin
ABC transporters in lipid homeostasis
spermatid differentiation
plasma membrane bounded cell projection assembly
Cell-extracellular matrix interactions
Hedgehog 'off' state
Complement cascade
regulation of hippo signaling
response to mechanical stimulus
peptidyl-serine phosphorylation
GeneRatio
0.25
0.20
0.15
0.10
0
1
2
3
4
-log(adj P-value)
C
Geometric Mean TPM - IMD AMPs
7500
5000
2500
0
Condition
D
Condition
AttA
AttB
AttC
AttD
CecA1
CecA2
Def
DptA
DptB
Dro
6
4
2
0
-2
-4
-6
log2FC over Ctrl
E
Survival Probability (%)
100
75
50
25
0
0 1 2 3 4 5 6 7
Days Post Injection
Condition
PBS
Efae Low Dose
Efae High Dose
Genotype
OrR
Imd[10191]
F
Survival Probability (%)
100
75
50
25
0
0 1 2 3 4 5 6 7
Days Post Re-Injection
Condition
PBS/PBS
Mock-Primed
Efae-Primed
Genotype
OrR
Imd[10191]

**Supplementary Figure 2: Additional analysis for fat body RNA-seq**

**A).** Heatmap of $log_2$FC over non-injected controls of whole-body, core immune response genes from Troha, et al. 2018.
Only a subset of the core genes were ubiquitously expressed across all conditions assayed for in this study. The differences are likey due to distinctions in time point and tissue. **B).** GO term enrichment from fat body Mock-Primed-specific, up-regulated genes. **C).** Geometric means of TPMs of IMD-dominant AMPs (as identified in FlyBase Gene Group FBgg0001194: Imd Signaling Pathway) across collected fat body samples (Mock-Primed vs *Efae*-Primed; Welch's t-Test: $p = 0.6562$). **D).** Heatmap of $log_2$FC over non-injected controls of IMD-dominant AMPs across collected fat body samples **E).** Single-injection survival comparison between OrR and *imd*-mutant flies **F).** Double-injection survival comparison between OrR and *imd*-mutant flies. Data from **D & E** are the same as in **Figure 3G&H**, replotted for comparison.
