## Supplementary Figure 3 for "Drosophila immune priming to Enterococcus faecalis relies on immune tolerance rather than resistance"

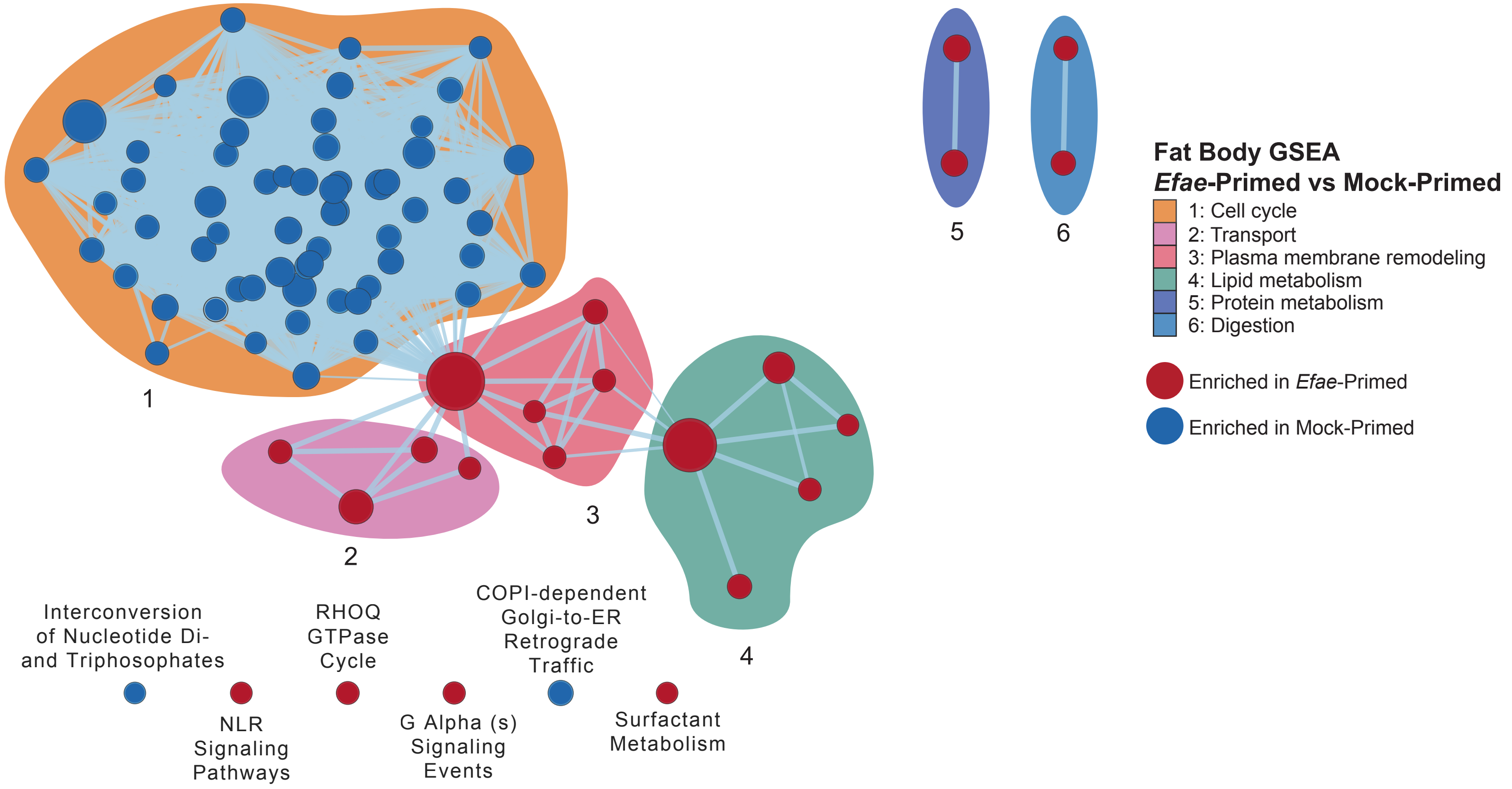


**Supplementary Figure 3: GSEA for Efae-primed versus Mock-primed fat bodies**

This visualization represents relationships between statistically significant terms (FDR < 0.05), manually curated with clusters that summarize the relationships between terms. Full results are found in **Supplementary Table 3**.
