## Supplementary Figure 4 for "Drosophila immune priming to Enterococcus faecalis relies on immune tolerance rather than resistance"

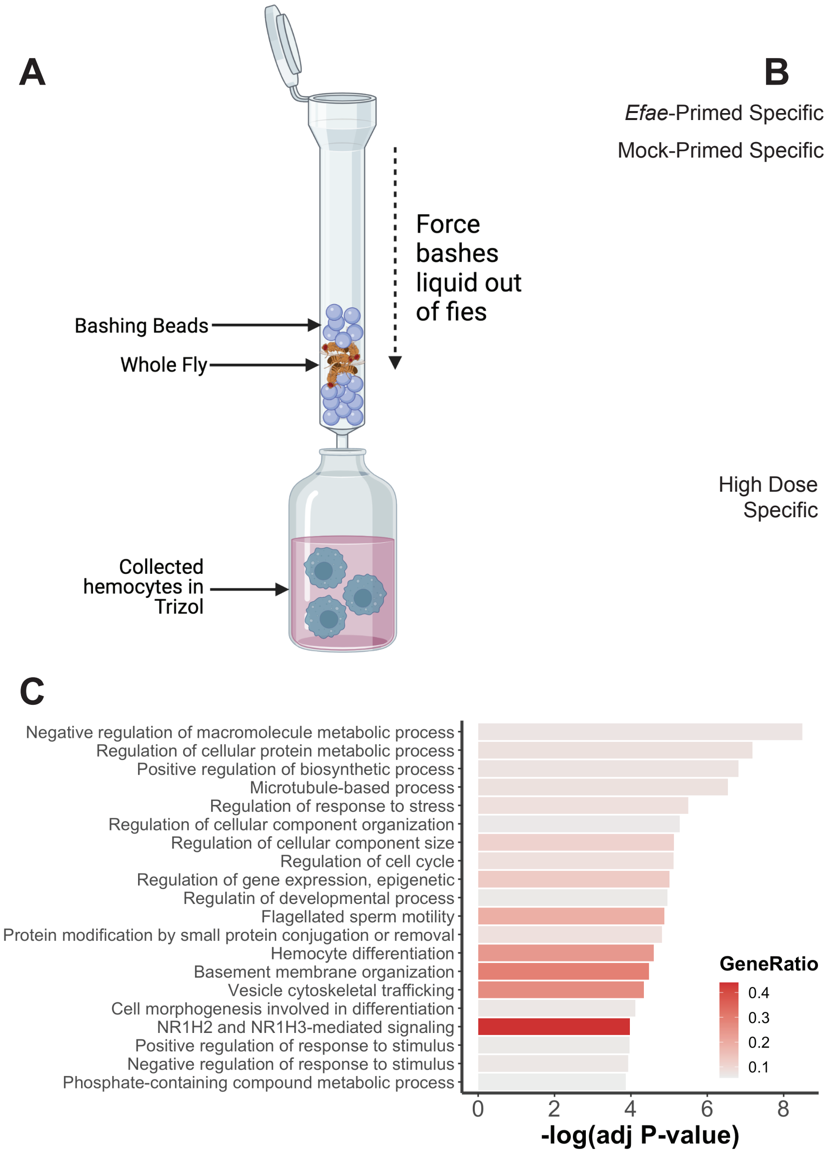


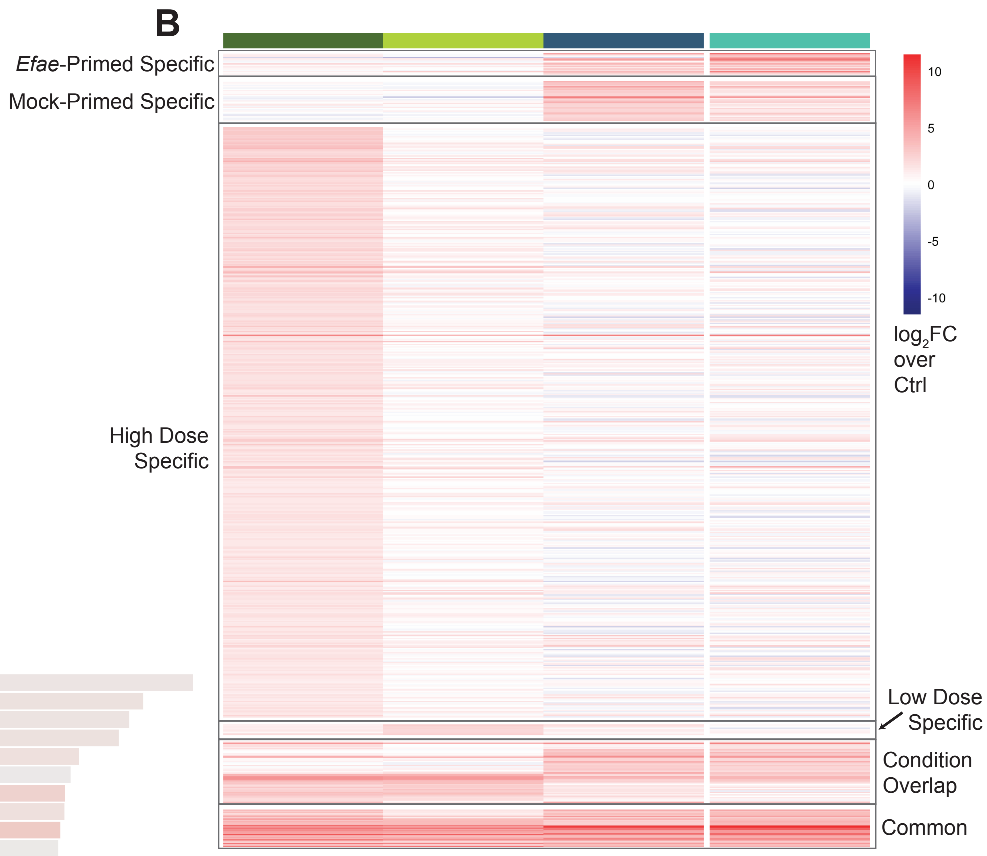


D

| FlybaseID | Name | Symbol |
|---|---|---|
| FBgn0027584 | Uncharacterized | CG4757 |
| FBgn0032639 | Uncharacterized | CG18563 |
| FBgn0283461 | Drosomycin | Drs |
| FBgn0032638 | Serine protease homolog 93 | SPH93 |
| FBgn0034407 | Diptericin B | DptB |
| FBgn0034511 | GNBP-like 3 | GNBP-like3 |
| FBgn0034328 | Bomanin Bicipital 1 | BomBc1 |
| FBgn0034331 | Bomanin Bicipital 2 | BomBc2 |
| FBgn0265577 | Inducted by Infection | IBIN |
| Fbgn0034329 | Bomanin Short 1 | BomS1 |
| Fbgn0262881 | Metchnikowin-like | Mtkl |

**Supplementary Figure 4: Additional data for hemocyte RNA-seq**

**A).** Schematic diagram of hemocyte RNA extraction. **B).** Significantly up-regulated genes as corresponding to conditions in **Fig 4A** (scale: $log_2$FC over age-matched controls) **C).** GO term enrichment from hemocyte *Efae* Hi Dose-specific, up-regulated genes. **D).** Overlap of up-regulated core genes (4-condition overlap in venn diagram) between hemocytes and fat bodies.
