## Supplementary Figure 5 for "Drosophila immune priming to Enterococcus faecalis relies on immune tolerance rather than resistance"

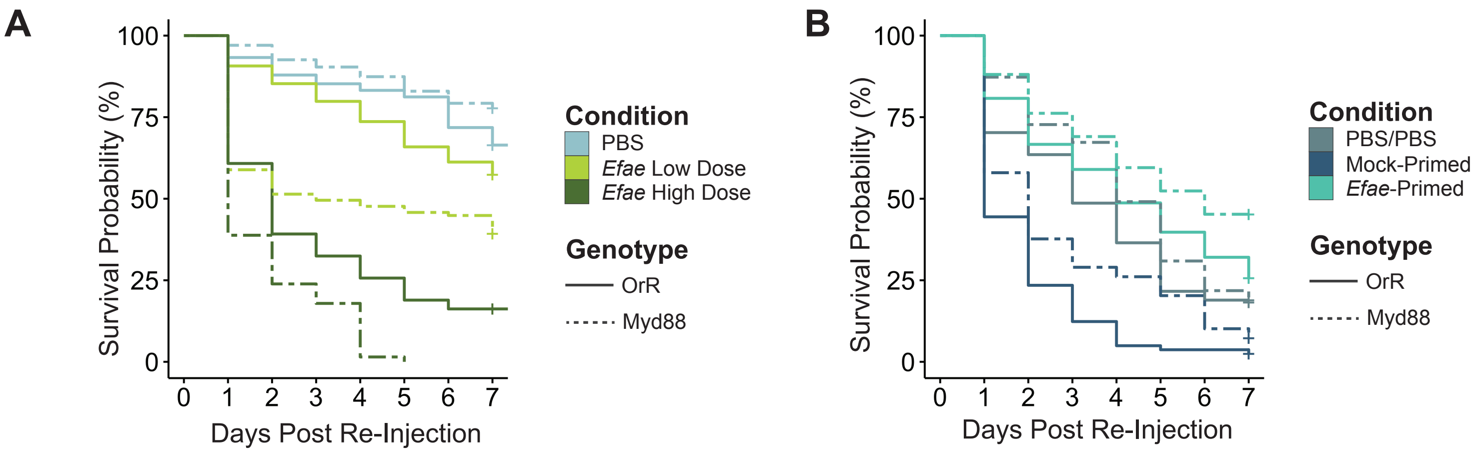


**Supplementary Figure 5: Additional data for immune loitering RNA-seq**

**A).** Single-injection survival comparison between OrR and *Myd88*-mutant flies **B).** Double-injection survival comparison between OrR and *Myd88*-mutant flies. Data are the same as in **Figure 5 D&E**, replotted for comparison.
