## Supplementary Figure 6 for "Drosophila immune priming to Enterococcus faecalis relies on immune tolerance rather than resistance"

A

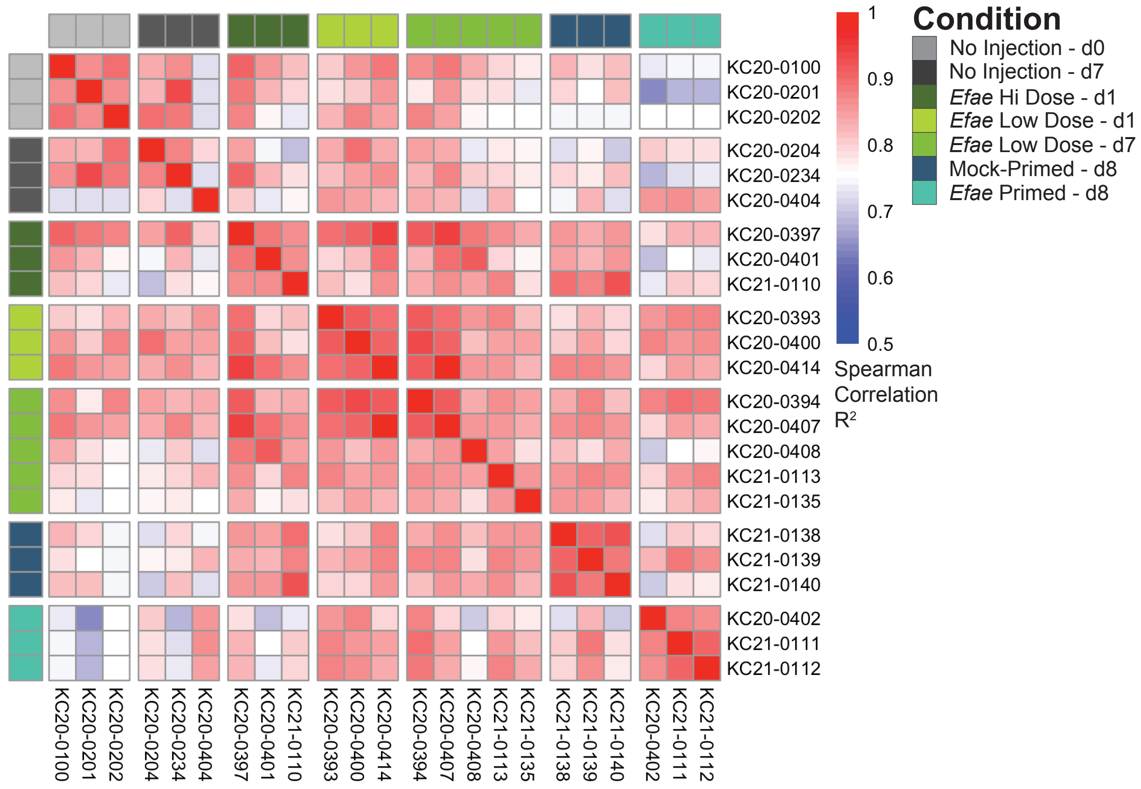


B

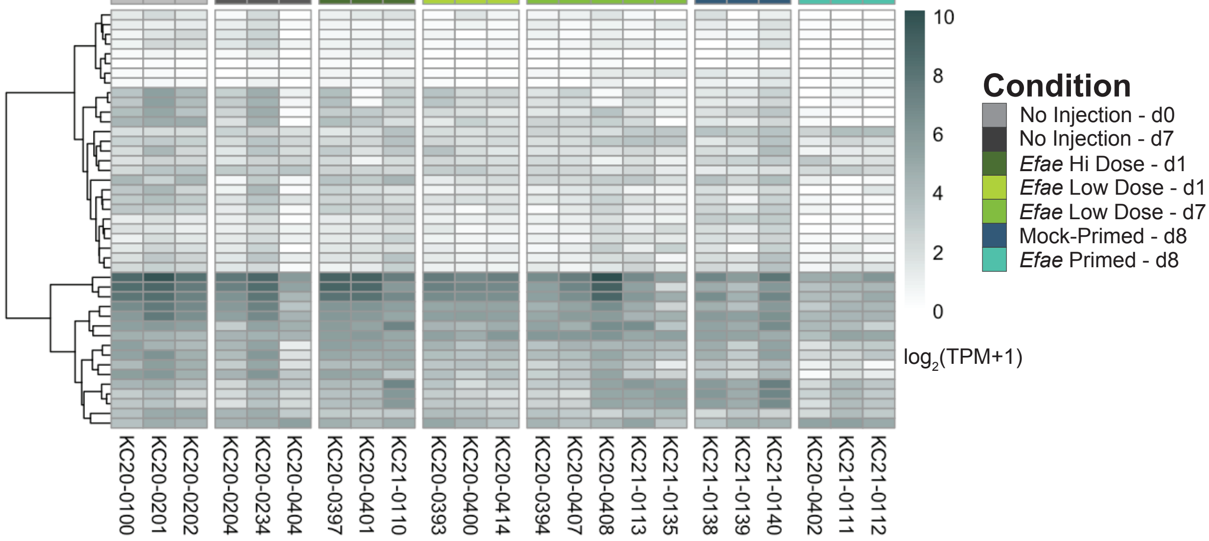


**Supplementary Figure 6: Quality control of fat body RNA-seq libraries**

**A).** Spearman correlation heatmap of fat body RNA-seq libraries. Values are $R^2$ spearman correlation values. **B).** Expression of sperm motility genes in fat body RNA-seq libraries. Values are $\log_2$(TPM+1).
