## Supplementary Figure 7 for "Drosophila immune priming to Enterococcus faecalis relies on immune tolerance rather than resistance"

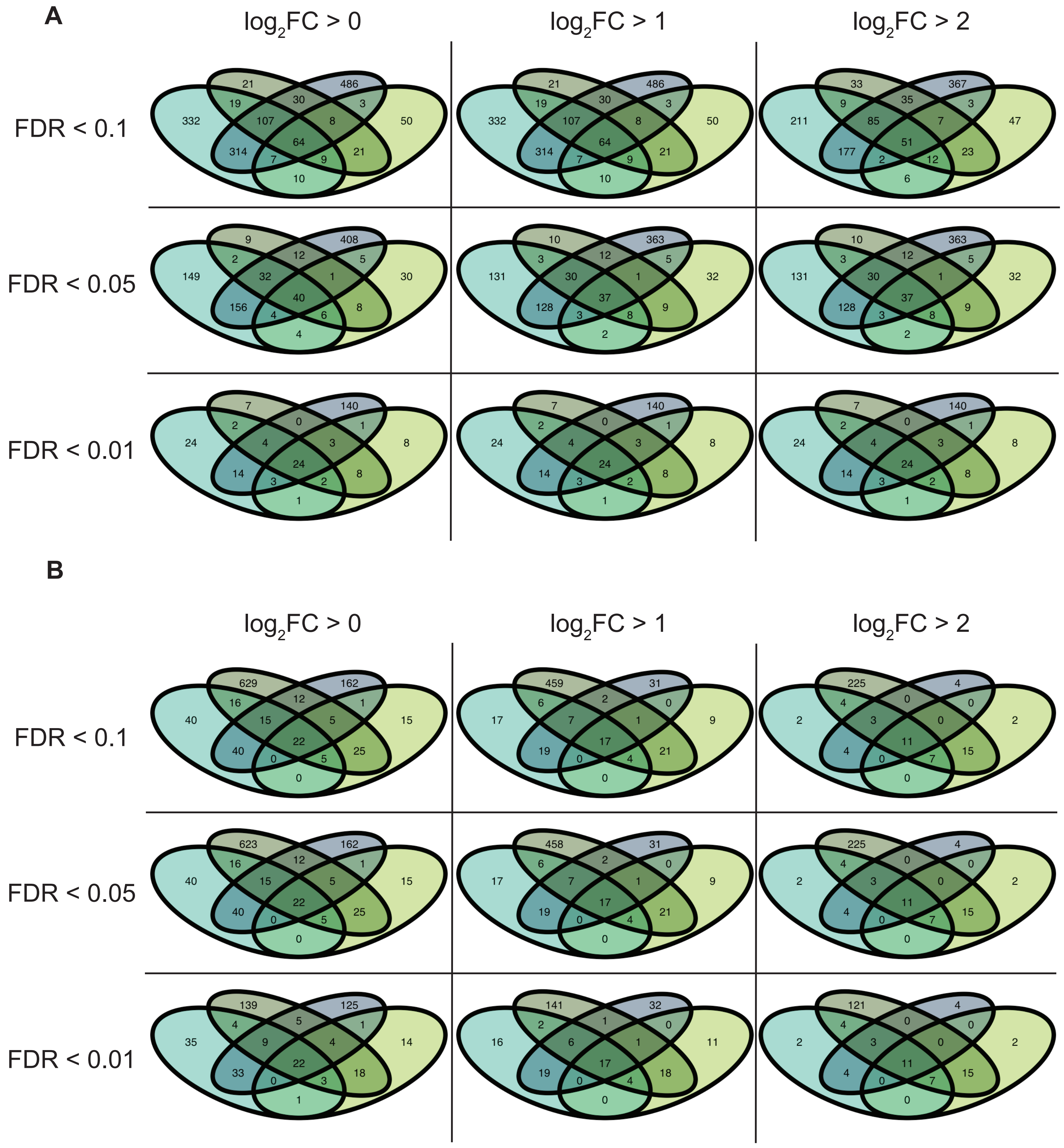


**Supplementary Figure 7: Modulation of significance and fold-change cutoffs in differential analysis**

**A).** Overlap analysis between 24-hour RNA-seq in fat bodies when changing fold-change cut-offs along the y-axis ($\log_2$FC>0, $\log_2$FC>1, logFC>2) and significance cutoffs along the x-axis (FDR<0.1, FDR<0.05, FDR<0.01). **B).** Same analysis as A, with hemocytes.
